# The development trends of antineoplastics targeting PD-1/PD-L1 based on scientometrics and patentometrics

**DOI:** 10.1101/2020.05.01.072041

**Authors:** Ting Zhang, Juan Chen, Yan Lu, Zhaolian Ouyang

## Abstract

**Background:** This paper aims to show the scientific research and technological development trends of antineoplastics targeting PD-1/PD-L1 based on scientometrics and patentometrics.

**Methodology/Principal Findings:** Publications and patents related to antineoplastics targeting PD-1/PD-L1were searched and collected from the Web of Science (WoS) and the Derwent Innovation Index (DII) respectively. Totally, 11244 publications and 5501 patents were obtained. The publications were analyzed from the annual number, the top countries/regions and organizations to describe the scientific research trends in this field. The patents were analyzed from the annual number, the top priority countries and patent assignees to reveal the characteristics and status of technological development. As well as the identification of scientific research focus and technological development focus was based on the title and abstract of the publications and patents, using the freely available computer program VOSviewer for clustering and visualization analysis. The number of scientific publications and patent applications showed obvious increase of 29.84% and 33.46% in recent ten years (2009-2018), respectively. Results suggested that the most productive countries/regions publishing on antineoplastics targeting PD-1/PD-L1 were USA and China, and the top three productive organizations were all from USA, including Harvard University, VA Boston Healthcare System (VA BHS) and University Of California System. There were four scientific research focus: (1) immune escape mechanism, (2) biomarkers related to efficacy and prognosis, (3) immune-related adverse event, and (4) drug design and preparation, and five technological development focus: (1) testing methods and apparatus, (2) indications related to carcinoma, (3) biomarkers related to diagnosis and prognosis, (4) small molecule inhibitors, and (5) indications other than carcinoma.

**Conclusions/Significance:** The results of this study presents an overview of the characteristics of research status and trends of antineoplastics targeting PD-1/PD-L1, which could help readers broaden innovative ideas and discover new technological opportunities, and also serve as important indicators for government policymaking.

## Introduction

Cancer is one of the causes of death around the world, which threatens human health and quality of life seriously. The International Agency for Research on Cancer (IARC) released the latest report “Global cancer statistics 2018” ^[1]^, which showed that, there were an estimated 18.1 million new cancer cases worldwide, with a death toll of 9.6 million in 2018, which had risen sharply comparing to 14.1 million new cases and 8.2 million deaths in 2012. The global cancer burden has further increased. The Report showed that 1 in 5 men and 1 in 6 women worldwide would suffer from cancer, and 1 in 8 men and 1 in 11 women would die. The data showed that ^[2,3]^, almost half of the world’s new cancer cases and more than half of cancer deaths occurred in Asia in 2018. Among them, China, as a populous country, was a large part of the cancer incidence and death in Asia. The incidence of cancer in China was increasing year by year and tending to be younger. In recent years, the World Health Organization (WHO) has proposed that, 40% of malignant tumors can be prevented, 40% of malignant tumors can be cured, and 20% of malignant tumors can be palliative, which could help cancer patients to live more comfortable. Early diagnosis and treatment can prolong survival time and improve quality of life. Antineoplastics have always been a hot spot in drug discovery, especially tumor immunotherapy drugs have received globally widespread attention. In recent years, immunological checkpoint inhibitors targeting the programmed death receptor-1 (PD-1)/programmed death receptor-1 ligand (PD-L1) pathway have become a research focus in the field of cancer treatment, which could be the most promising tumor immunotherapy strategy, due to its long-lasting effect and wide applicability. PD-1 is an important immunosuppressive molecule that is closely related to cancer treatment. In 2012, Suzanne LT and Julie R found that PD-1 related antibodies had a good effect in non-small cell lung cancer (NSCLC), melanoma and kidney cancer ^[4,5]^, which was the first time to be confirmed that PD-1 related antibodies had been shown to be effective in cancer. PD-1 is a negative costimulatory signaling molecule, and PD-L1 is its ligand. The PD-1/PD-L1 signaling pathway is one of a group of immunostimulatory molecules that plays a key role in immune activation and tolerance. A series of subsequent evidences suggest that PD-1 targeted immunomodulation plays an important role in anti-tumor therapy ^[6–10]^. In 2013, Science magazine ranked tumor immunotherapy as the top of the top ten breakthroughs ^[11,12]^, which once again made immunotherapy the focus of cancer treatment, and the research on PD-1/PD-L1 provided new strategies for anti-tumor treatment. PD-1/PD-L1 monoclonal antibody has become the standard treatment for a variety of malignant tumors, such as malignant melanoma, non-small cell lung cancer (NSCLC), Hodgkin’s lymphoma, renal cell carcinoma, etc. Since 2014, the US Food and Drug Administration (FDA) has approved five immunological checkpoint inhibitors targeting the PD-1/PD-L1 pathway, including two PD-1 monoclonal antibodies (Pembrolizumab and Nivolumab) and three PD-L1 monoclonal antibodies (Atezolizumab, Avelumab and Durvalumab), which have made breakthroughs in the clinical treatment of various tumors. These drugs have significantly prolonged the survival of many cancer patients, and some patients have completely resolved their condition. Merck’s Keytruda (Pembrolizumab) is the first PD-1 inhibitor approved by the FDA for the treatment of metastatic melanoma ^[13,14]^. A number of PD-1/PD-L1 monoclonal antibodies have been in clinical research all over the world, and antibody-based immunotherapy has become an important research area. PD-1/PD-L1 is a co-inhibitory molecule expressed on the surface of various tumors, which participates in the immune escape of tumor cells by down-regulating the anti-tumor activity of T cells. Blocking the PD-1/PD-L1 pathway has become the focus of tumor immunotherapy. These drugs show a broad spectrum of cancer treatment, and they are expected to achieve a huge breakthrough in cancer treatment in the future. The treatment of PD-1/PD-L1 pathway has become a cancer immunotherapy. Antineoplastics targeting PD-1/PD-L1 develop rapidly, bringing hope to the majority of cancer patients. However, while focusing on their therapeutic effects, the limitations and side effects should also be clearly recognized. Exploring new ideas and directions will become a research focus in the future. The research on the antineoplastics targeting PD-1/PD-L1 is the future development trends, which has good application value and practical significance. With the accelerating pace of technology updates, the uncertainty of technological innovation is also increasing. It is helpful for technology forecasting by extracting effective information from massive data, and mining hidden inter-technology relationships.

As the most effective carrier of technical information, patents cover more than 90% of the latest technological information in the world, and the content is detailed and accurate. Patent analysis is the theory and method of obtaining intelligence from patent information, and it is the main method for analyzing the key technologies in information science. Through the patent analysis in a certain field, it can objectively reflect the overall situation and development trend of technology ^[15–18]^, which could effectively and accurately give the information about the relevant technological innovation and development. Antineoplastics targeting PD-1/PD-L1 are developing rapidly and are the focus of international competition. The patent analysis of antineoplastics targeting PD-1/PD-L1 could help us grasp the technological innovation trend and clarify the direction of technological innovation. This study could provide information support for the research of cancer drug therapy in China, and also provide a new research perspective for technological development.

### Ethics Statement

This study is about the scientometrics and patentometrics of antineoplastics targeting PD-1/PD-L1, and the subject investigated are the academic publications and patents. It does not involve human subject research and animal research, and no patient records/information and clinical records are included. Therefore, no ethics issue is involved in this research.

## Materials and Methods

### Materials

The publications and the patents were both from the database of Thomson Reuters, with two-way connection, which could organically link the basic research results and technological application ones. Then it made it possible to understand the relationship between basic research results and market application prospects, analyze the competitive situation globally, and accelerate the mutual promotion and transformation of knowledge innovation and technology.

The publications related to “PD-1” and “PD-L1” were searched and collected from the Web of Science (WoS) Database of Thomson Reuters, using search strategy based on topic in May, 2019. There were two subject terms of “PD-1” and “PD-L1”, which were retrieved in title and abstract. The search strategy was as follows: Topic “PD-1” OR Topic “PD-L1”, and then refined by document types as “articles” and languages as “English”, and then 11244 ones were obtained.

The patents were collected from the Derwent Innovation Index (DII) Database of Thomson Reuters using search strategy based on subject terms of “PD-1” and “PD-L1”, which were also retrieved in title and abstract. Then, 5495 patents were obtained.

The publication and patent data were imported into Thomson Data Analyzer 3.0 (TDA, Thomson Reuters Co., New York, NY, USA) for authority control, and imported into the freely available computer program VOSviewer (version 1.6.3; www.vosviewer.com) for clustering and visualization analysis.

### Methods

Publications and patents are important outputs, which could reflect the situation of basic research, applied research and other aspects in one field, and also reveal the level of science and technology and international competitiveness of one country to a certain extent.

Scientometrics and patentometrics were employed to investigate the research activity and to reveal the current characteristics and status in the field of antineoplastics targeting PD-1/PD-L1. The scientometrics were based on the following descriptions of publication years, countries/regions, organizations and scientific research focus. The identification of scientific research focus were in view of co-word analysis of the title and abstract of the publications in this field. The patentometrics were in terms of application years, priority countries, patent assignees and technological development focus. The identification of technological development focus were also according to co-word analysis of the title and abstract of the patents in this field.

Scientometrics and patentometrics are primary methods in information science, which utilizes quantitative analysis to reveal the development situation in a given field on the basis of publications and patents, respectively. Scientometrics and patentometrics can provide a macroscopic overview of the given field for the researchers, and have been used in many scientific fields, e.g. oncology ^[15]^, neurosciences ^[19]^ and nanotechnology ^[20]^, etc. Through the scientometrics and patentometrics of the field of antineoplastics targeting PD-1/PD-L1, we can generally understand the main research direction and its development trend.

### The clustering and visualization by VOSviewer

VOSviewer, a freely available computer program, is developed for constructing and viewing bibliometric maps ^[21]^, has been widely used in many fields, such as computer and information ethics ^[22]^, physiology ^[23]^, 3D bioprinting ^[24]^, etc. In our research, the VOSviewer software constructed maps of scientific research focus and technological development focus, on account of the cooccurrence frequencies of the words from the title and abstract of the publications (2015-2019) and patents, respectively. The 2D maps in which the distance between any two words reflects the relatedness of the words as closely as possible. In general, the stronger the relation between two words, the smaller the distance between them in the map. Each word in the map also has a color. Colors are used to indicate the grouping or clustering of the words. Those words with the same color belong to the same cluster and tend to be more closely related than those with different colors. In other words, words with the same color tend to co-occur with each other more frequently than those with different colors. To obtain the visualization and the clustering of the words, the VOSviewer software uses two techniques, namely the VOS mapping technique and the VOS clustering technique, which share the same underlying principles, and together these techniques provide a unified framework for mapping and clustering ^[21,25]^. In the identification of scientific research focus and technological development focus, the map provide a visual representation by showing the relations between the words of the title and abstract of the publications (2015-2019) and patents in the field of t antineoplastics targeting PD-1/PD-L1, by means of using the VOSviewer software for bibliometric mapping ^[21]^. Those maps can be used to get an overview of the field of antineoplastics targeting PD-1/PD-L1 both in the scientific research and technological development aspects.

## Results and Discussions

### The scientific research trends of antineoplastics targeting PD-1/PD-L1 based on scientometrics

#### General analysis

Based on the above data sources and search strategy, 11244 articles were retrieved in all. Fig. 1 shows the growth of publications in the field of antineoplastics targeting PD-1/PD-L1. The year 2019 was not included due to that it was partial of the year (from January to May in 2019). The number of publications assigned annually increased from 1 in 1971 to 2391 in 2018, and showed an obvious increase from 2009, with an increase of 29.84% from 2009 to 2018, indicating that more and more attention has been paid on in recent years. An exponential regression was done based on the data from 2009 (the year in which the publication trend began to increase continuously) through 2018. The equation describing the data was y = 147.71e^0.2627x^ with a coefficient of determination R^2^ = 0.9507. If the publication increase continues at the same rate, the equation forecasts that 2656 documents will be published in 2019 and 7596 in 2023. Since the WOS database continues to include new journals, the actual numbers will be higher than the predicted values in 2019 and 2023.

**Fig 1.**
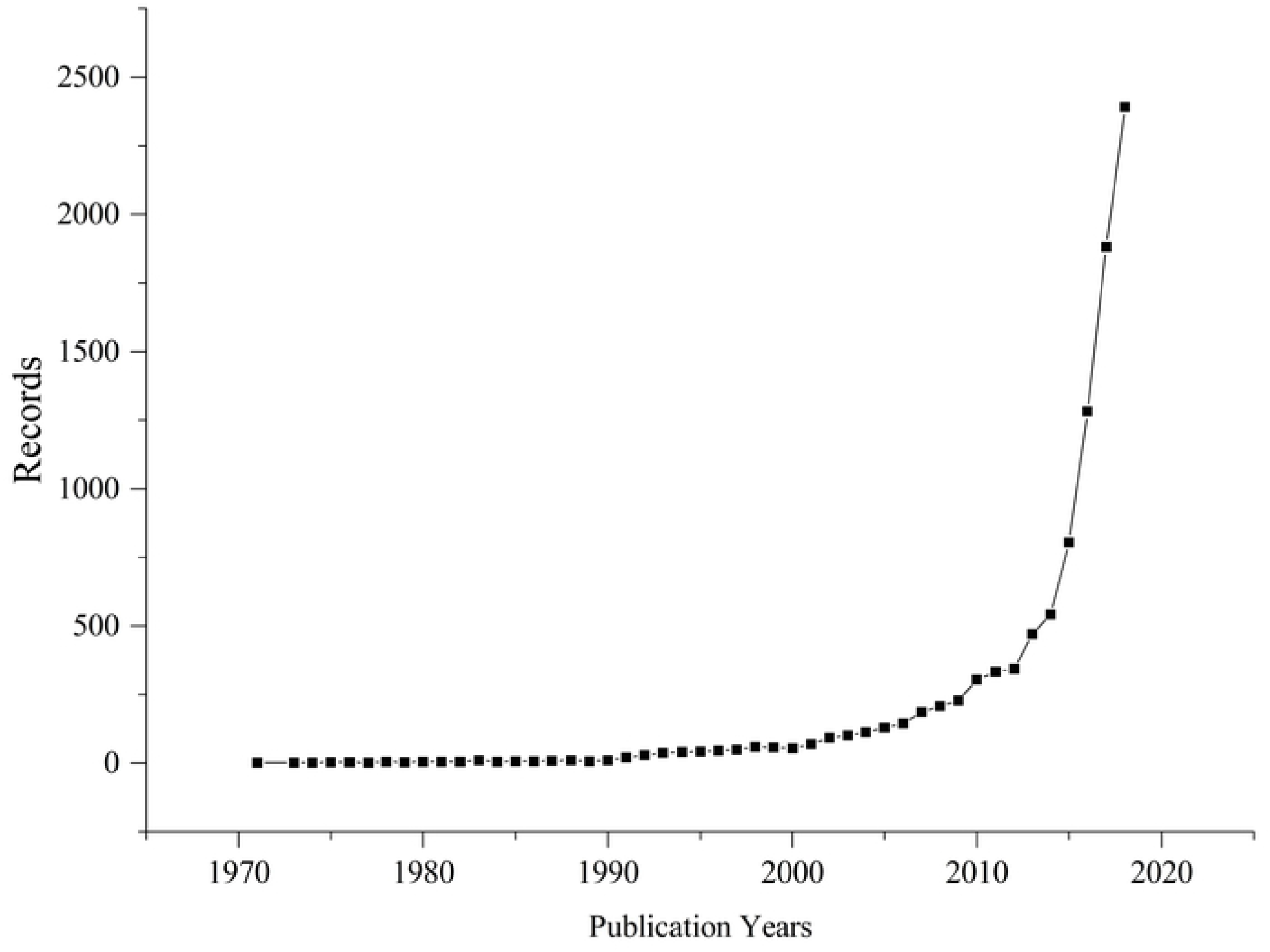
Annual changes in the numbers of the publications in the field of antineoplastics targeting PD-1/PD-L1.

#### Countries/Regions

A total of 113 countries/regions contributed to the field of antineoplastics targeting PD-1/PD-L1. The top ten prolific countries/regions were shown in Fig 2. USA had the greatest share of publications, followed by China and Japan. In particular, USA (4567 articles) published more than 40% of all the publications, over twice as many as articles as the following China (2226 articles). Similarly, China had close to twice numbers of articles as the following Japan (1230 articles). The total number of publications produced by these three countries was 8023 (71.35%), which constituted more than 70% of the global productivity in the field of antineoplastics targeting PD-1/PD-L1.

**Fig 2.**
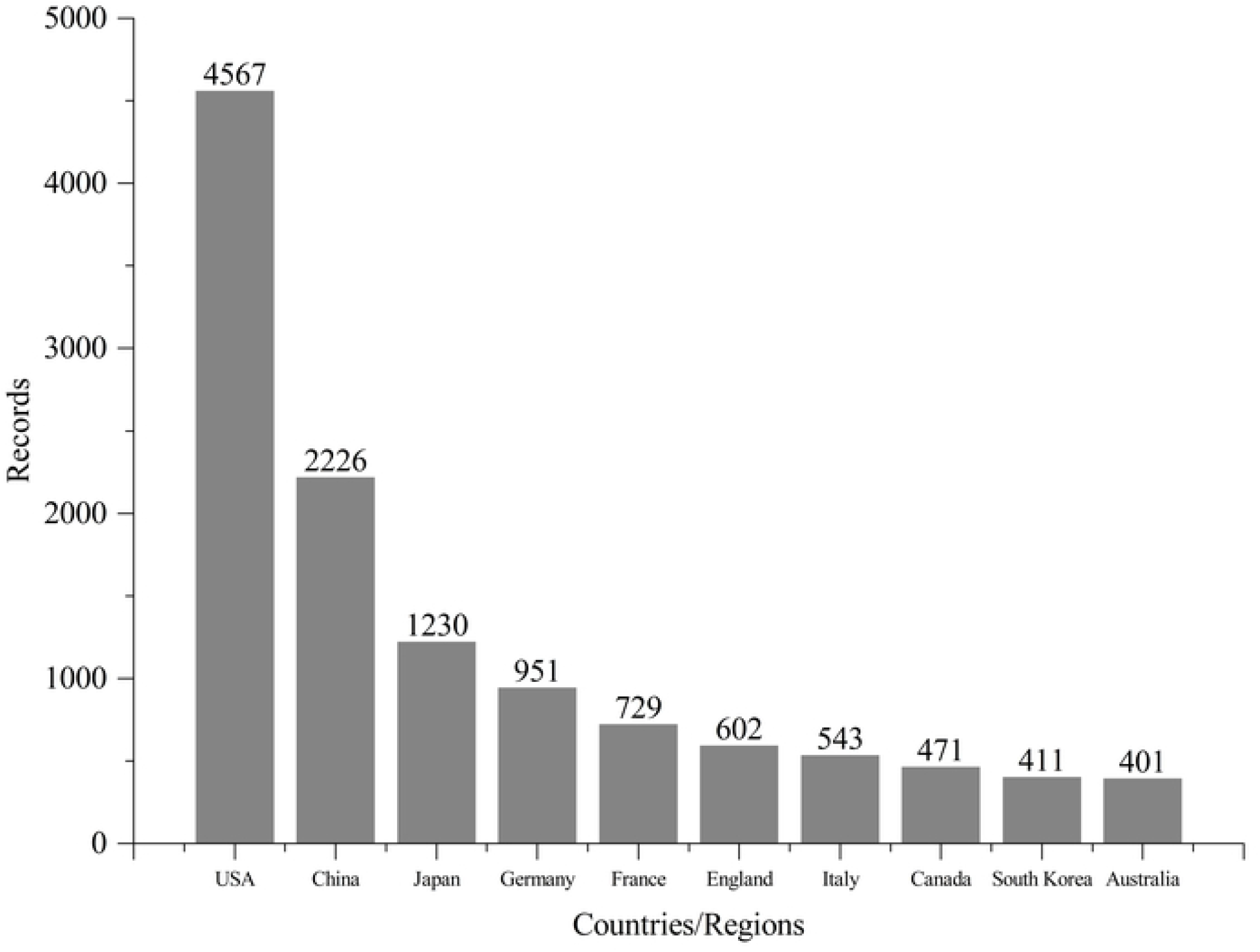
The top ten prolific countries/regions in the field of antineoplastics targeting PD-1/PD-L1.

In the recent five years, the number of publications has exceeded half of the total number of publications in this field, so the recent five years have been chosen to reflect the trend of publications in the top 10 countries/regions. Fig. 3 showed the annual changing trends of the top 10 countries/regions from 2015 to 2018. Also, the year 2019 was not included due to the incomplete data (from January to May in 2019), which could not represent the final trend. The results indicated that the publications of USA and China increased most rapidly in recent years. Both the total number and the annual number of publications in recent years in China were in the second place. However, there is still a wide gap between China and USA.

**Fig 3.**
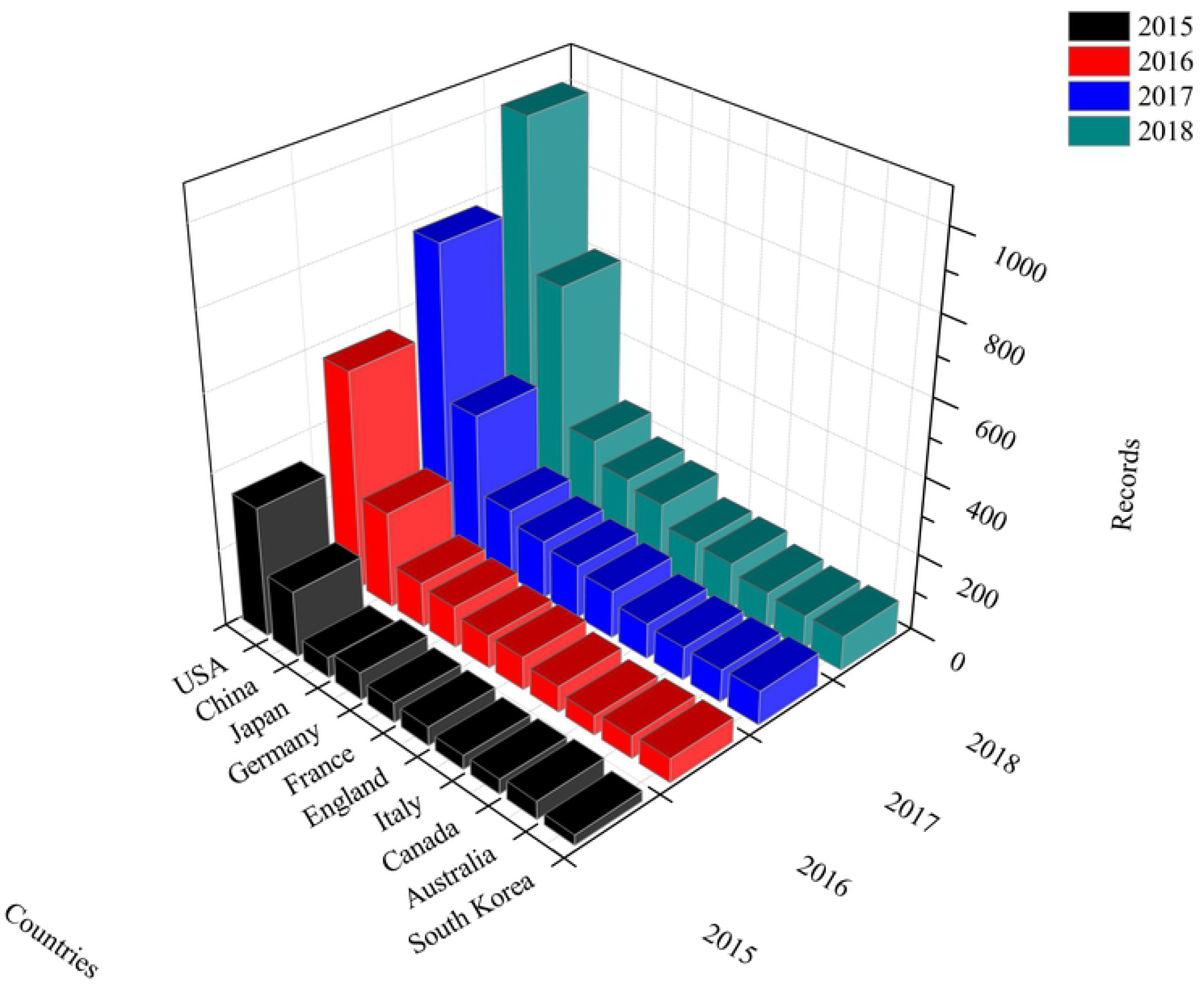
The annual changing trends of the top 10 countries/regions in the field of antineoplastics targeting PD-1/PD-L1 from 2015 to 2018.

#### Organizations

The top 20 organizations were shown in Fig. 4, which were highly correlated with the top 10 countries/regions. More than half of them were American, with 14 ones. There were three institutions in France entering the top 20, and one institution in China, one in Germany and one in England entered the top 20, respectively. The top three organizations were Harvard University, VA Boston Healthcare System (VA BHS) and University Of California System, with publications of 763, 590 and 469, respectively. The only one Chinese organization in top 20 was Chinese academy of sciences, with 216 publications.

**Fig 4.**
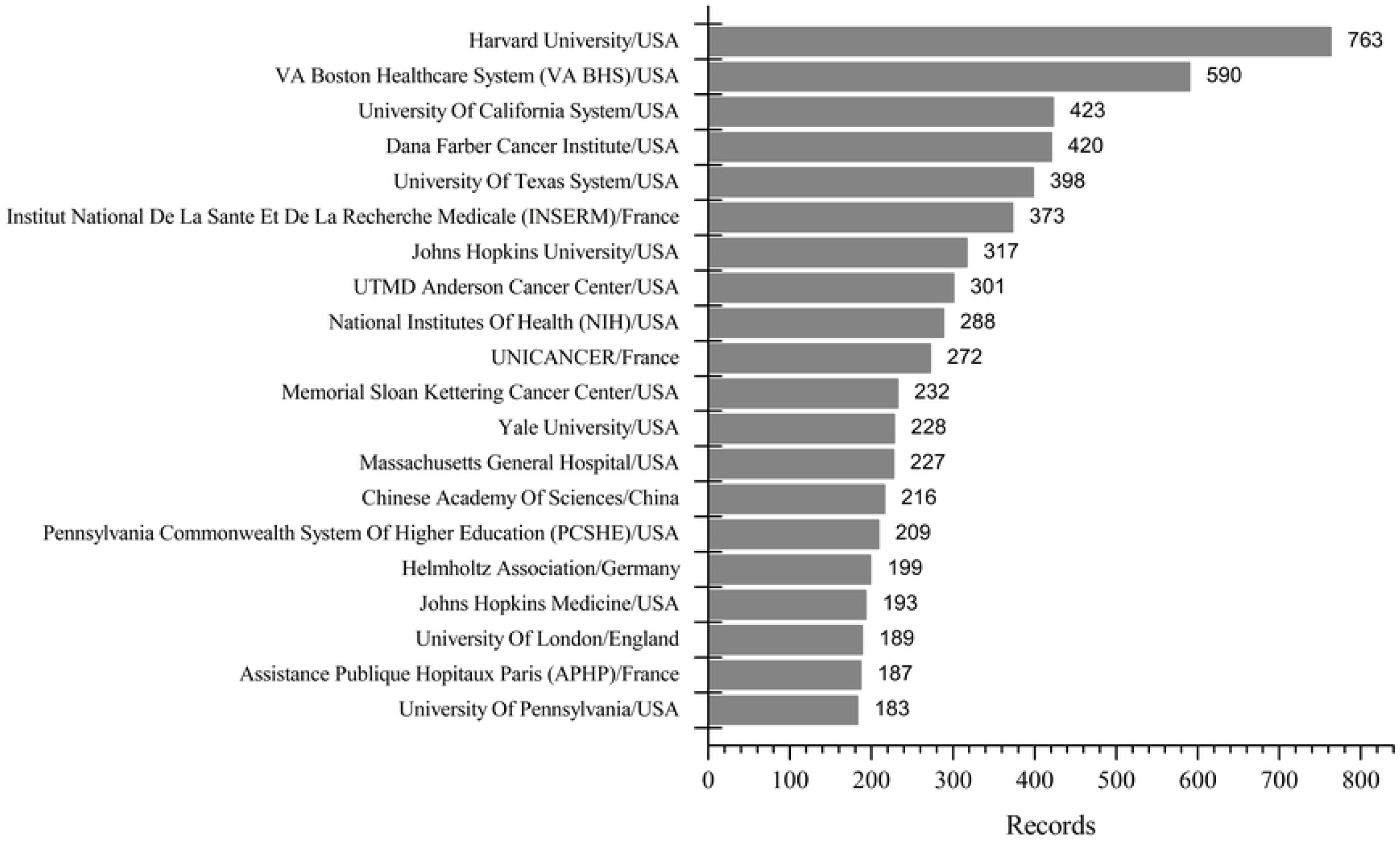
The top 20 organizations in the field of antineoplastics targeting PD-1/PD-L1 from 2015 to 2018.

#### Scientific Research Focus

There were 11244 publications related to antineoplastics targeting PD-1/PD-L1, including 7506 ones in recent five years (2015-2019). The titles and abstracts of the 7506 ones were chosen to identify the scientific research focus, which were imported to VOSviewer for clustering and visualization. Fig 5 showed the clusters of co-word matrix made by VOSviewer, and there were divided into four clusters, indicating that the scientific research focus of related to antineoplastics targeting PD-1/PD-L1 were varied. Each cluster is one scientific research focus, the four focus were as follows: (1) immune escape mechanism, (2) biomarkers related to efficacy and prognosis, (3) immune-related adverse event, and (4) drug design and preparation (Fig 5A). Those words associated with each focus were listed in Table 1, and each cluster only listed the top ten high frequency words due to the limited space. Fig 5B showed the temporal development of the scientific research focus related to antineoplastics targeting PD-1/PD-L1 in recent five years, which indicated that the four focus were concentrated in 2016 and 2017, and the focus of efficacy and prognosis, and immune related adverse event were newer than immune escape mechanism, and drug design and preparation in recent five years, relatively, which also indicated that the focus of biomarkers related to efficacy and prognosis, and immune-related adverse event had received more attention recently.

**Table 1.**
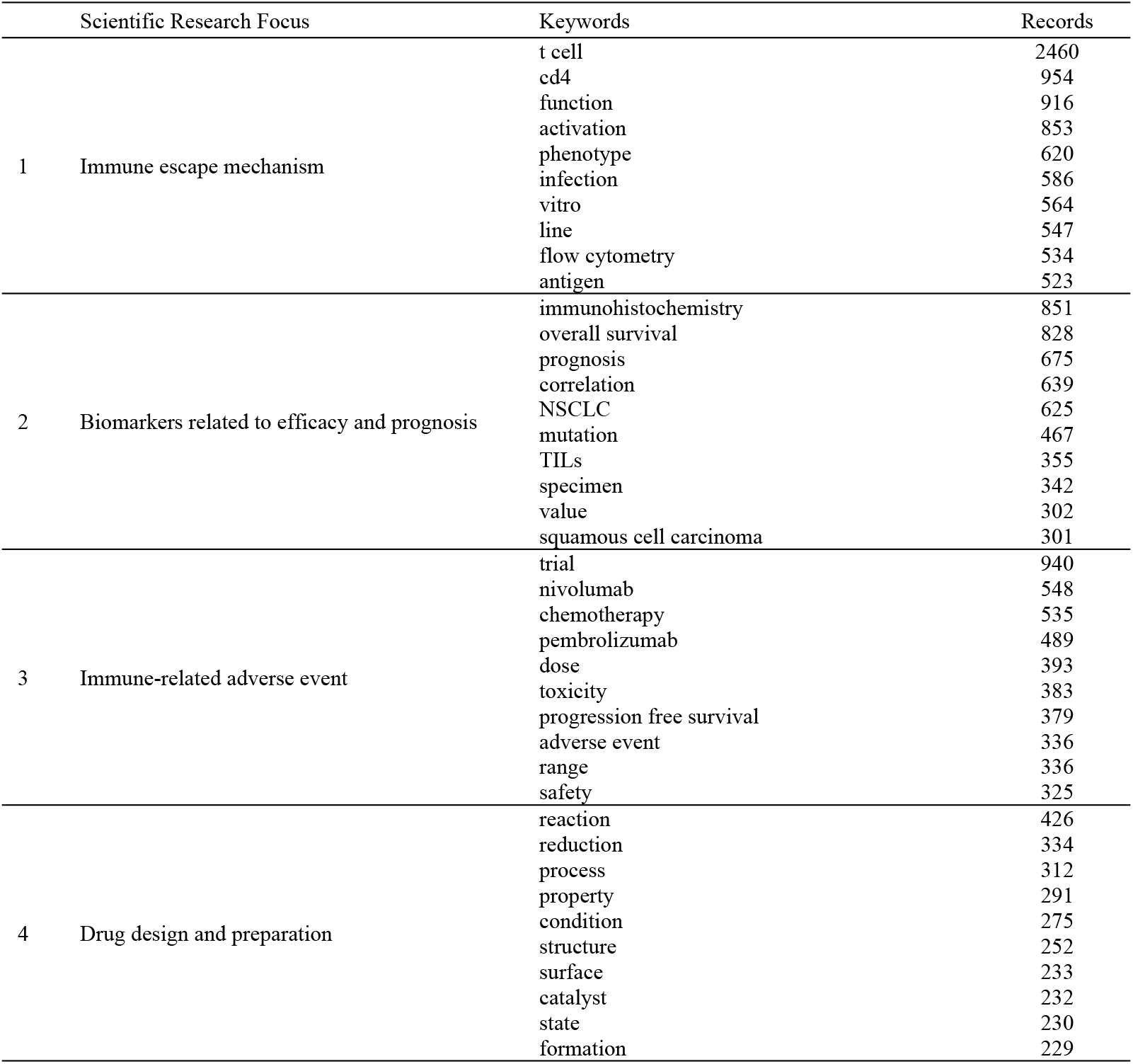
The scientific research focus related to antineoplastics targeting PD-1/PD-L1.

**Fig 5.**
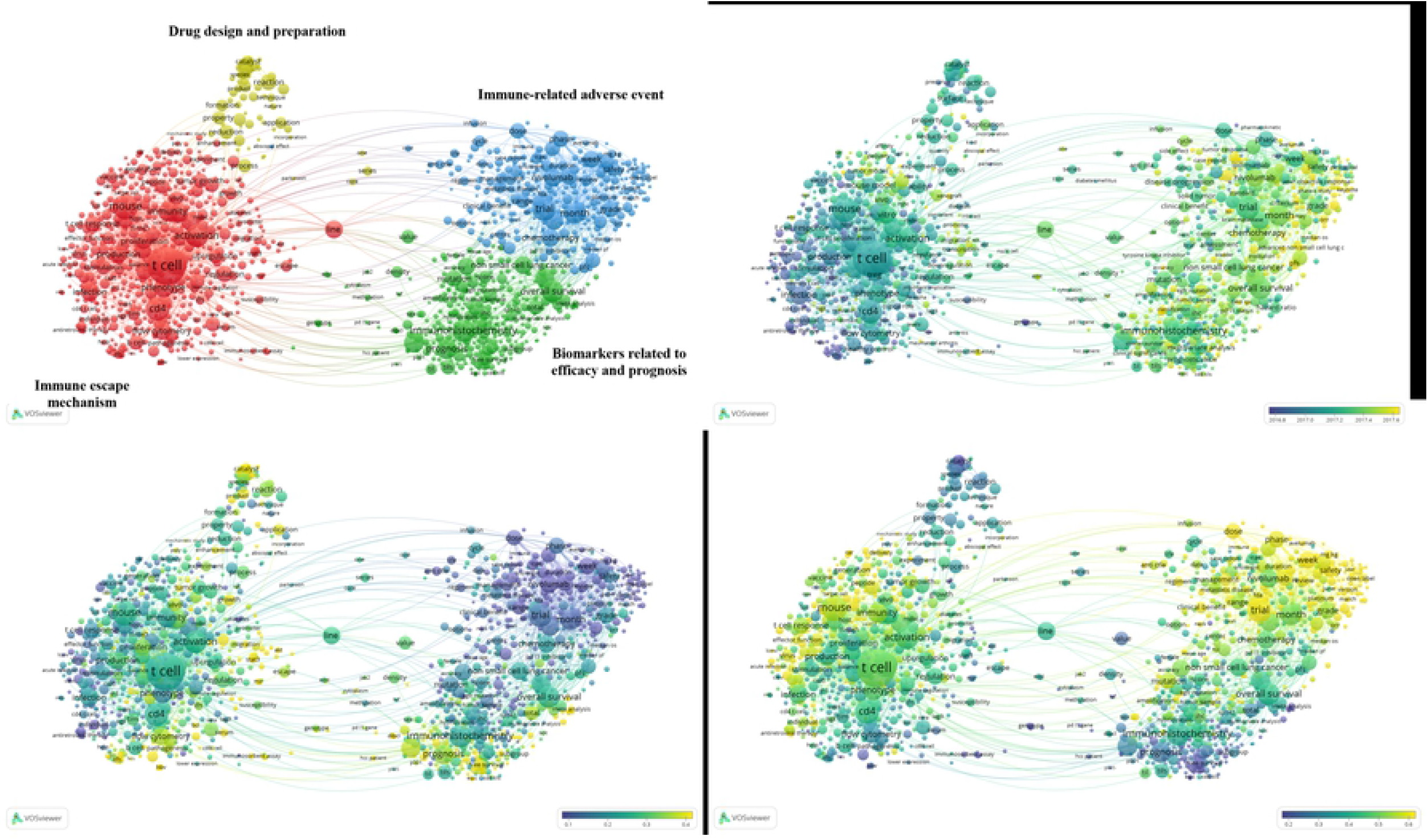
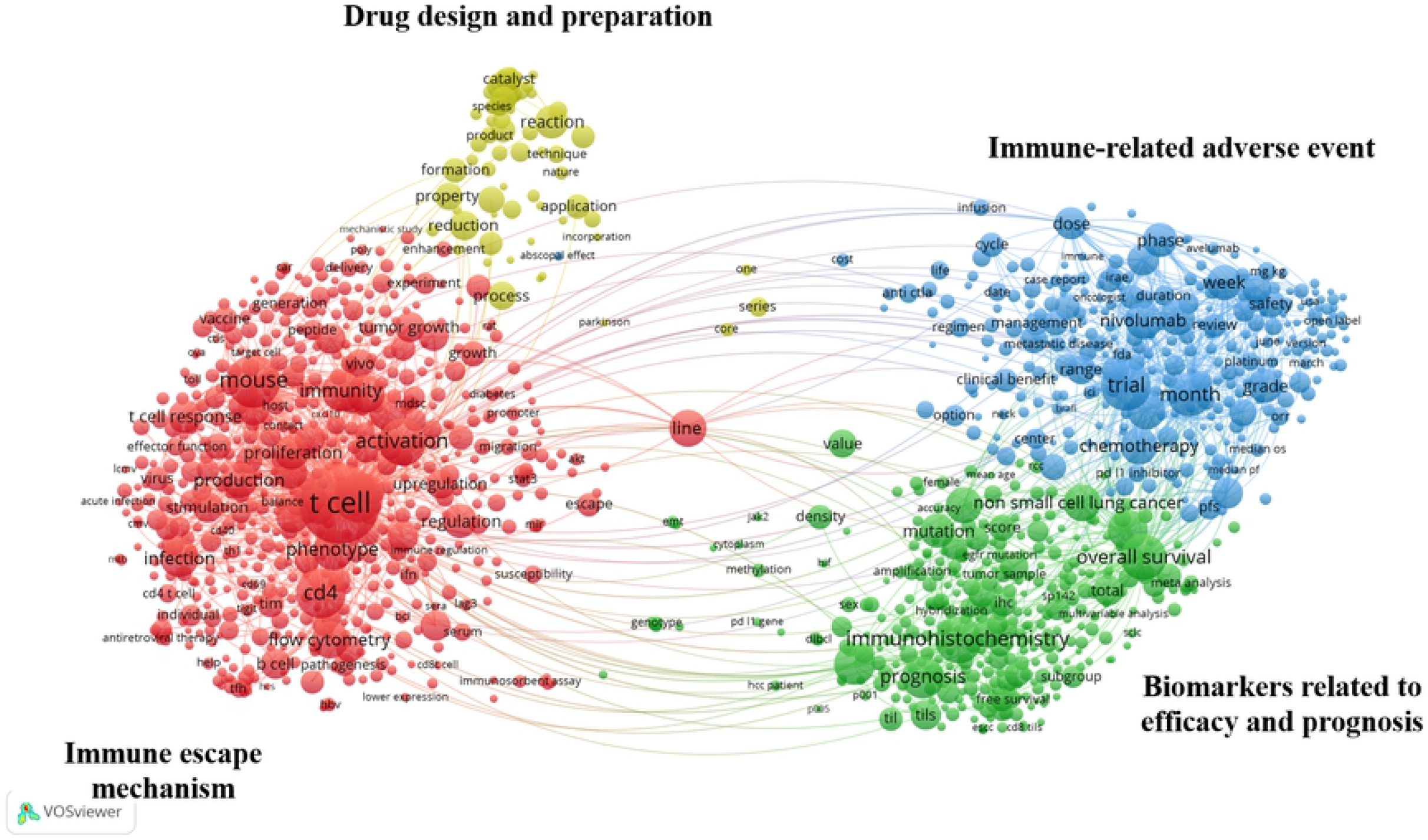
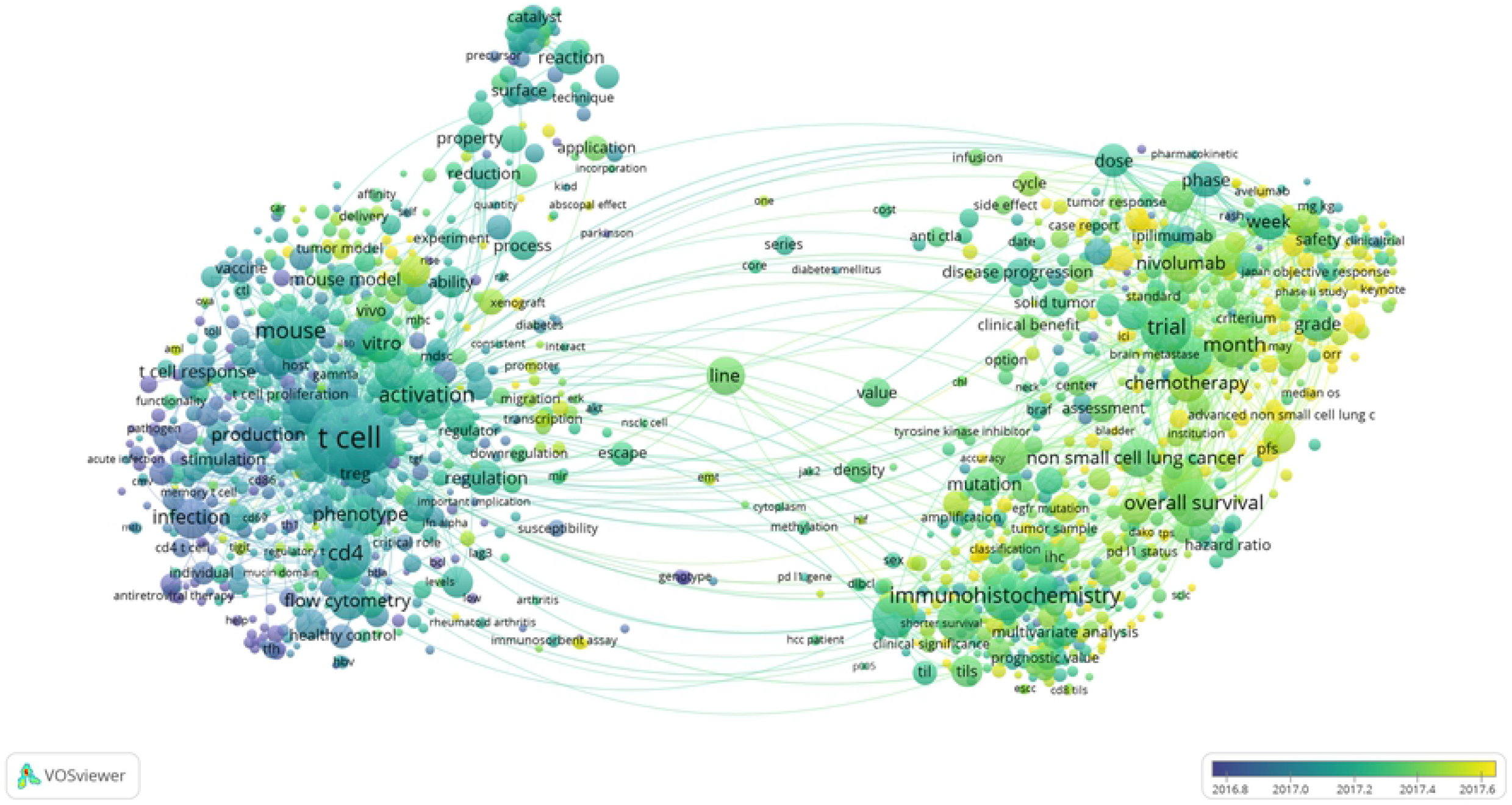
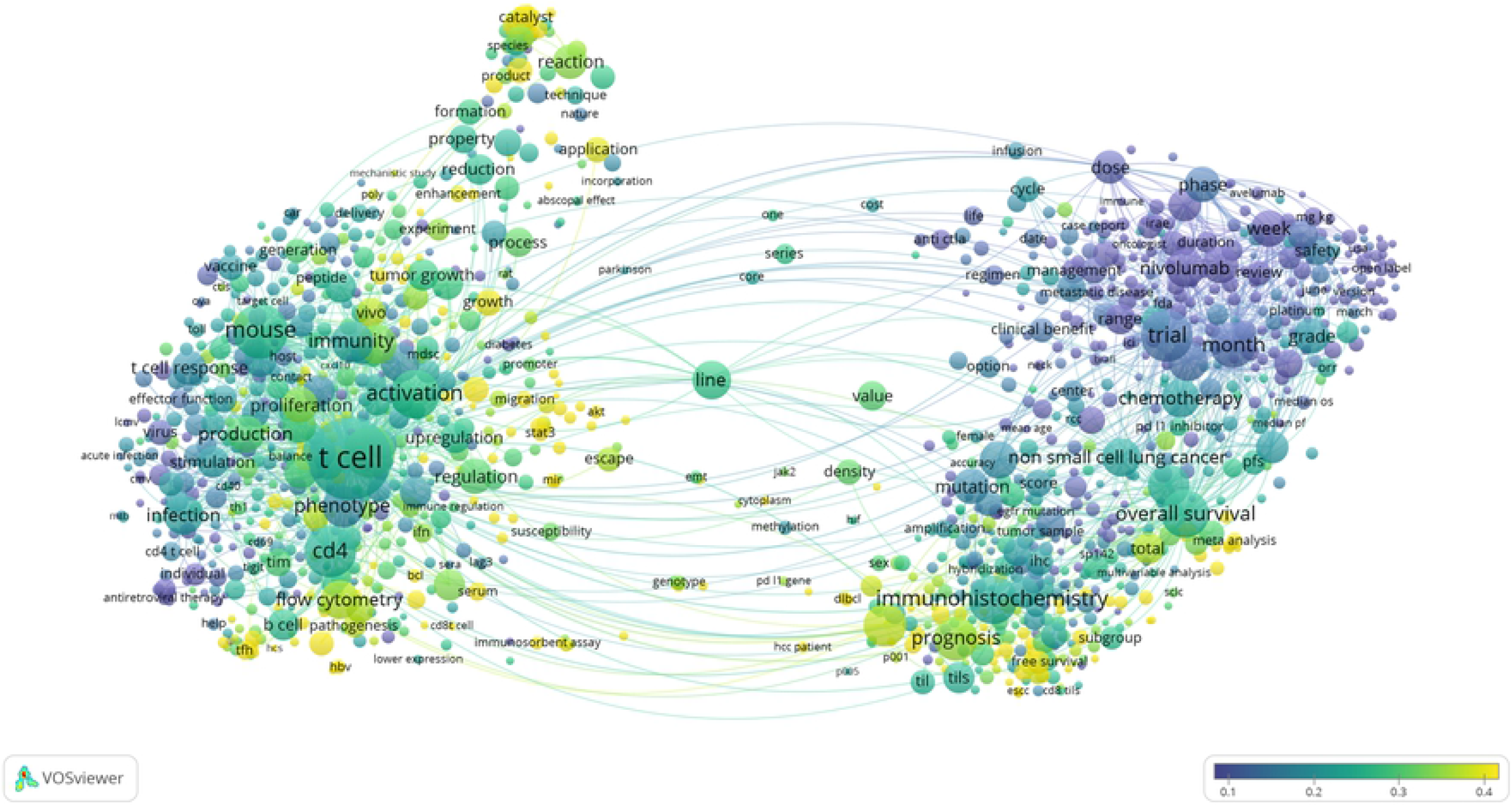
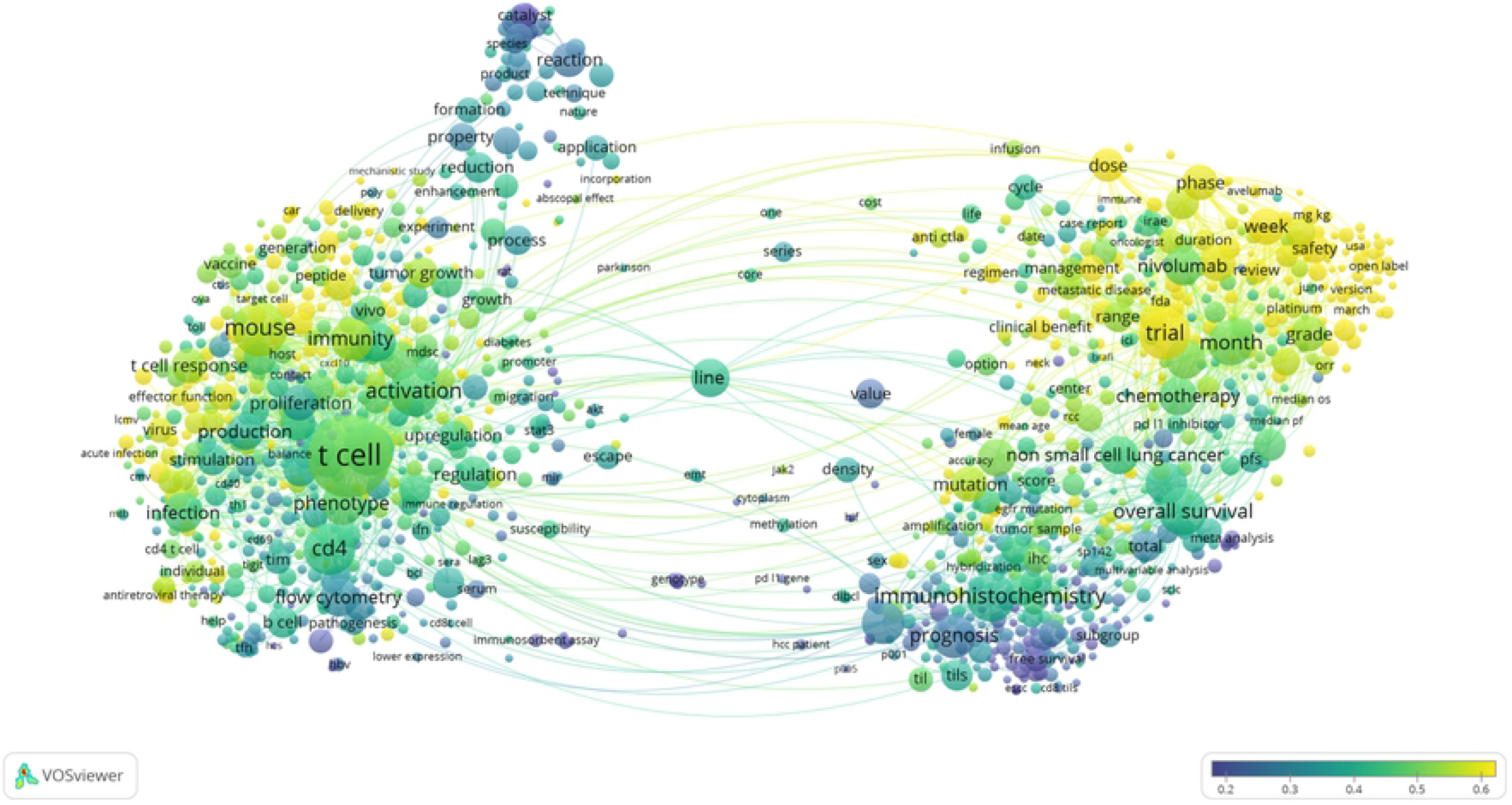
The scientific research focus related to antineoplastics targeting PD-1/PD-L1. **(A)** The visualization of the scientific research focus related to antineoplastics targeting PD-1/PD-L1 based on the VOSviewer. **(B)** The temporal development of the scientific research focus related to antineoplastics targeting PD-1/PD-L1 in recent five years. The color is from purple to yellow, the closer to purple, the focus is older, and the closer to yellow, the focus is newer. **(C)** The scientific research focus of China related to antineoplastics targeting PD-1/PD-L1. Colors from purple to yellow indicate that the concern of focus is low to high, with the value from 0.1 to 0.4. The closer to the purple (0.1), the focus is lower concerned, and the closer to the yellow (0.4), the focus is more concerned. **(D)** The scientific research focus of USA related to antineoplastics targeting PD-1/PD-L1. Colors from purple to yellow indicate that the concern of focus is low to high, with the value from 0.2 to 0.6. The closer to the purple (0.2), the focus is lower concerned, and the closer to the yellow (0.6), the focus is more concerned.

##### Biomarkers related to efficacy and prognosis

Immune checkpoint inhibitors have been shown to be effective in many malignancies, but the efficacy of different types of tumors is very different, and even patients with the same type of cancer respond differently ^[26–30]^. Therefore, finding effective biomarkers has always been crucial, which can screen those patients who could benefit from immunotherapy, and monitor the efficacy and prognosis of immunotherapy. At present, the commonly used biomarkers in clinical include programmed cell death 1 ligand 1(PD-L1) ^[31,32]^, tumor mutation burden (TMB) ^[33,34]^, microsatellite instability-high (MSI-H) and mismatch repair deficiency (dMMR) ^[35,36]^. However, most of these biomarkers need to be detected in tumor tissues, with the problems difficult acquisition and the low prediction efficiency. A recent clinical study identified the dMMR / MSI-H as a biomarker for the clinical response of PD-1 inhibitors, and FDA has accelerated approval of pembrolizumab for adults and pediatric patients with unresectable or metastatic dMMR or MEI-H solid tumors based on it ^[37]^. That was the first FDA-approved treatment option based on biomarkers rather than tumor types. It is an urgent problem to find the convenient and reliable biomarkers related to efficacy and prognosis in the field of immunotherapy. By exploring the biomarkers related to efficacy and prognosis of immunotherapy, it is possible to gain a deeper understanding of the interaction between the tumor and the immune system, and further determine the correct individualized treatment plan to make patients get more effective treatment. Identifying patients who could benefit from treatment, developing reliable biomarkers, and minimizing side effects from treatment will remain the research priorities.

##### Immune-related adverse events

Immune checkpoint inhibitors offer new hope for tumor patients, but also accompanying by new toxicity and disease. Immune-related adverse events (irAEs) refers to the imbalance of autoimmune tolerance caused by the enhancement of T cell tumor immunity, showing inflammatory response when normal tissues are involved ^[38]^. The commonly affected are the gastrointestinal tract, the skin, and the endocrine system ^[39]^, and the skin toxicity is the most common immune-related adverse event in the treatment of immune checkpoint inhibitors among them ^[40]^. In addition, there are pulmonary toxicity, renal toxicity, neurotoxicity, ocular toxicity, and rheumatic toxicity, which are relatively rare ^[39]^. Although pulmonary toxicity is uncommon, its toxicity occurs rapidly and can even cause death ^[41–43]^. Also, neurotoxicity is rare, but can be very serious and life-threatening ^[44,45]^. The toxic and side effects of immune checkpoint inhibitors are directly related to the excessive activation of the immune system, which are obviously different from those of chemotherapy and small molecule inhibitors. The side effects associated with chemotherapy and small molecule inhibitors are usually relieved by themselves after drug withdrawal, while immune-related adverse events are unique, and their toxicity may delay the onset and continue for months after drug withdrawal. It is usually impossible to solve immune-related adverse events simply by stopping medication or measuring adjustments, and it often requires the intervention of hormones or other drugs to inhibit the immune response ^[39,46]^. Early identification and intervention of the side effects of immune checkpoint inhibitor therapy are necessary conditions to reduce the incidence of adverse reactions and mortality, and also are essential to prevent long-term morbidity.

Fig 5C and Fig 5D showed the different scientific research focus between China and USA. It seems that China’s focus are more concerned on biomarkers related to efficacy and prognosis, immune escape mechanism, and drug design and preparation than immune-related adverse event. However, American focus are more concerned on immune-related adverse event, and immune escape mechanism than biomarkers related to efficacy and prognosis, and drug design and preparation. The scientific research focus of the two countries is a little different, and China has one more focus of drug design and preparation. However, in terms of the values of the two same focus in the two countries, the two focus of USA (near 0.6) is more concerned than those of China (near 0.4).

### The technological development trends of antineoplastics targeting PD-1/PD-L1 based on patentometrics

#### General analysis

Patents of antineoplastics targeting PD-1/PD-L1were collected from the Derwent Innovation Index (DII) Database of Thomson Reuters, and 5501 patents were obtained. The first patent application in this field was in 1990, with three ones. Then, the number of patents increased year by year, especially in recent years, showing an obvious increase from 2009. Fig. 6 showed the growth of patent applications in the field of antineoplastics targeting PD-1/PD-L1. The year 2019 was not included due to the lag period from patent application to publication, which could not reflect the real trends. The number of patent applications assigned annually increased from 3 in 1990 to 1988 in 2018, and showed an obvious increase from 2014, with an increase of 50.74% from 2014 to 2018, indicating that more and more attention had been paid on in recent years. A polynomial regression was done based on the data from 2010 (the year in which patent application began to increase continuously) through 2018. The equation describing the data was y = 42.463x^2^ - 179.63x + 306.38 with a coefficient of determination R^2^ = 0.976. If the patent application increase continues at the same rate, the equation forecasts that there will be 2855 patent applications in 2019 and 6553 in 2023.

**Fig 6.**
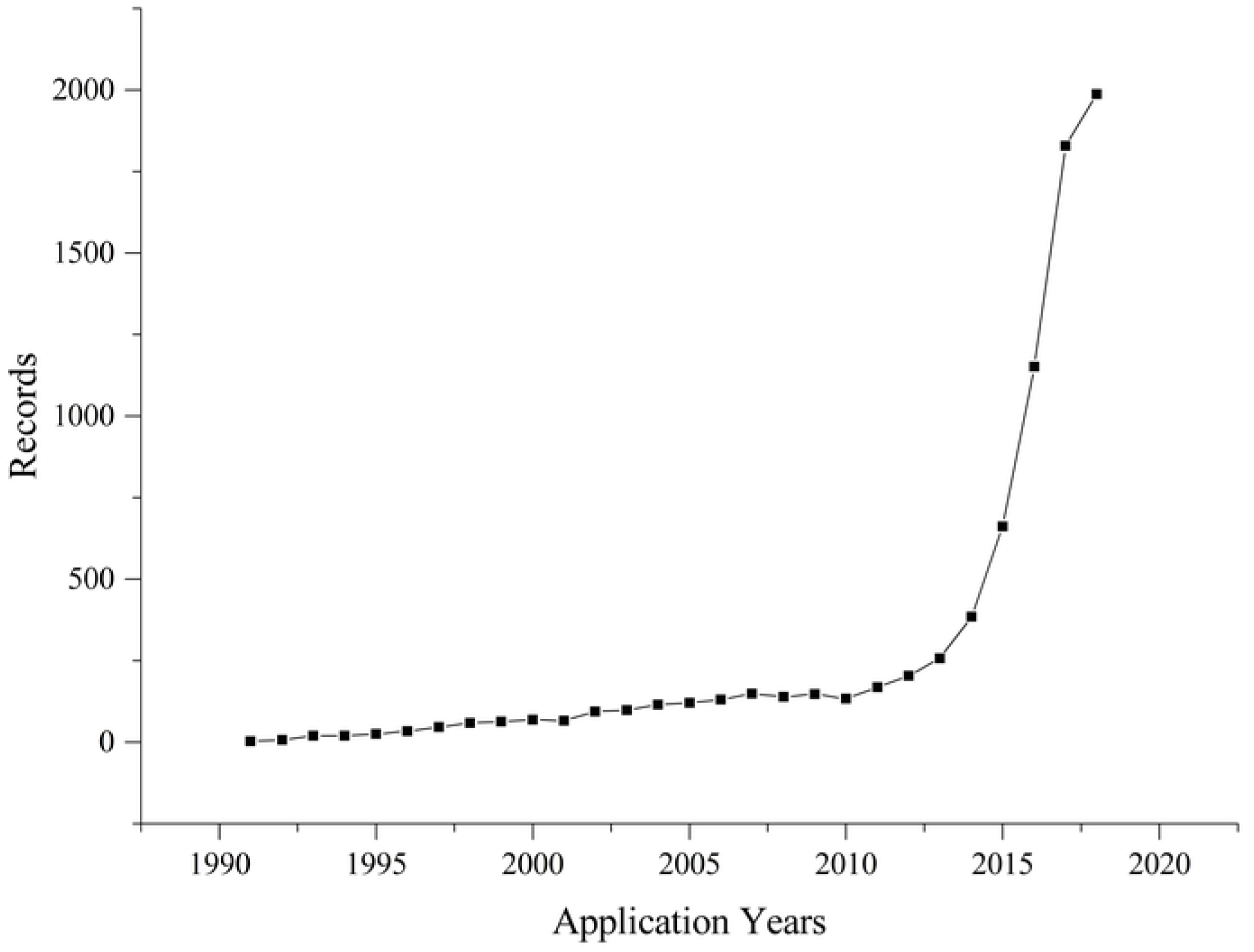
Annual changes in the numbers of the patent applications in the field of antineoplastics targeting PD-1/PD-L1.

#### Priority Countries

The top 10 priority countries were shown in Fig 7. The priority country with the largest number of patents was the United States (US, 2803), followed by China (CN, 1257) and Canada (CA, 1081). The United States’ patent applications were more than half of the worldwide, much than twice of the following China and Canada. The patent applications of Japan was 826, and the other priority countries was less than 500.

**Fig 7.**
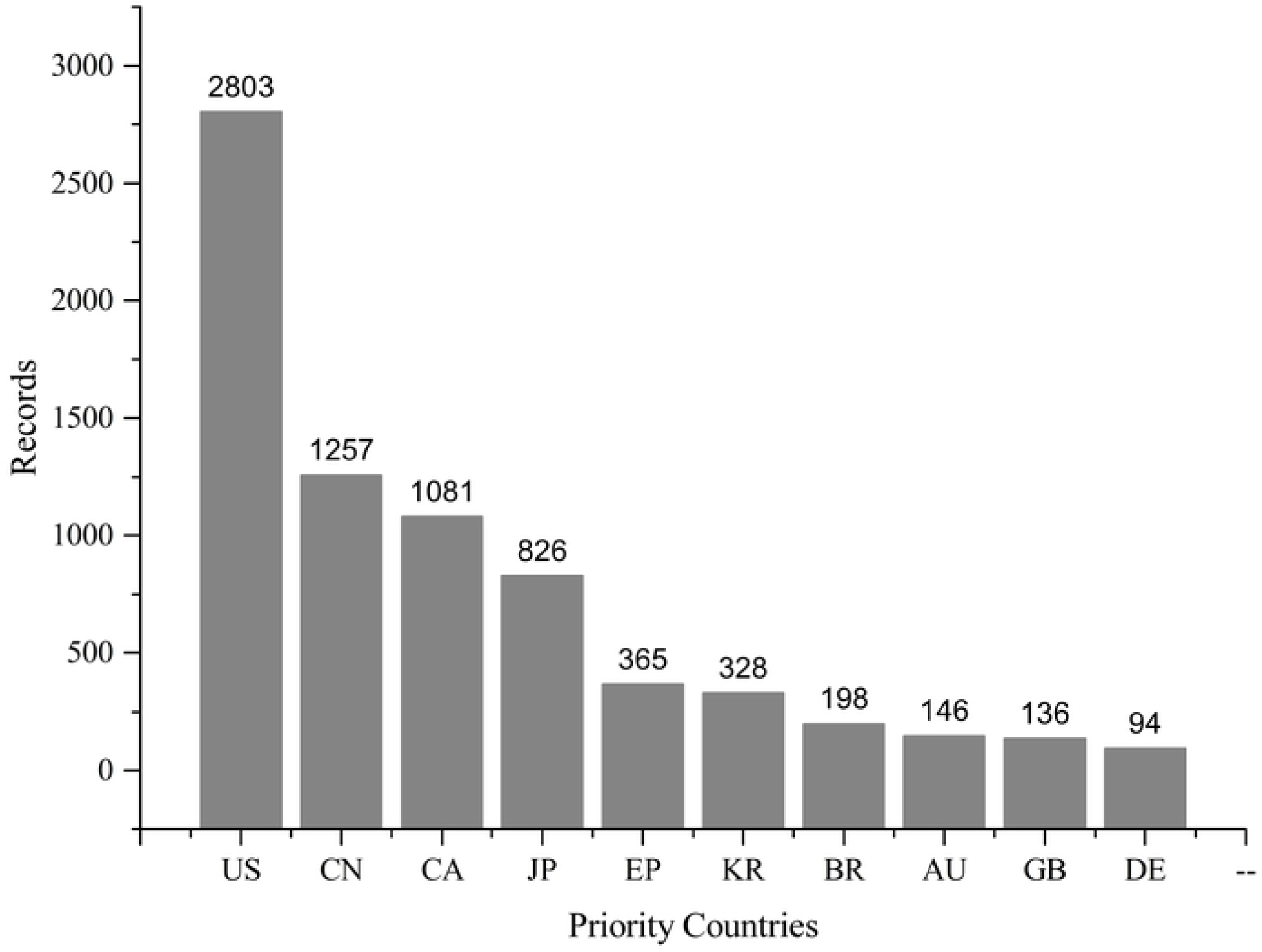
The top 10 priority countries in the field of antineoplastics targeting PD-1/PD-L1.

Fig. 8 showed the annual number of patent applications related to antineoplastics targeting PD-1/PD-L1 by priority countries, from 2009 to 2018. It depicted the rising trend of global patent applications on antineoplastics targeting PD-1/PD-L1, mainly in the United States, China and Canada. The annual number of patent applications in the United States ranked first in the world each year in recent ten years. Although the total number of patent applications in Canada was less than in China, the number per year was above China. Until 2018, the number of applications in China exceeded that of Canada, ranking second in the world.

**Fig 8.**
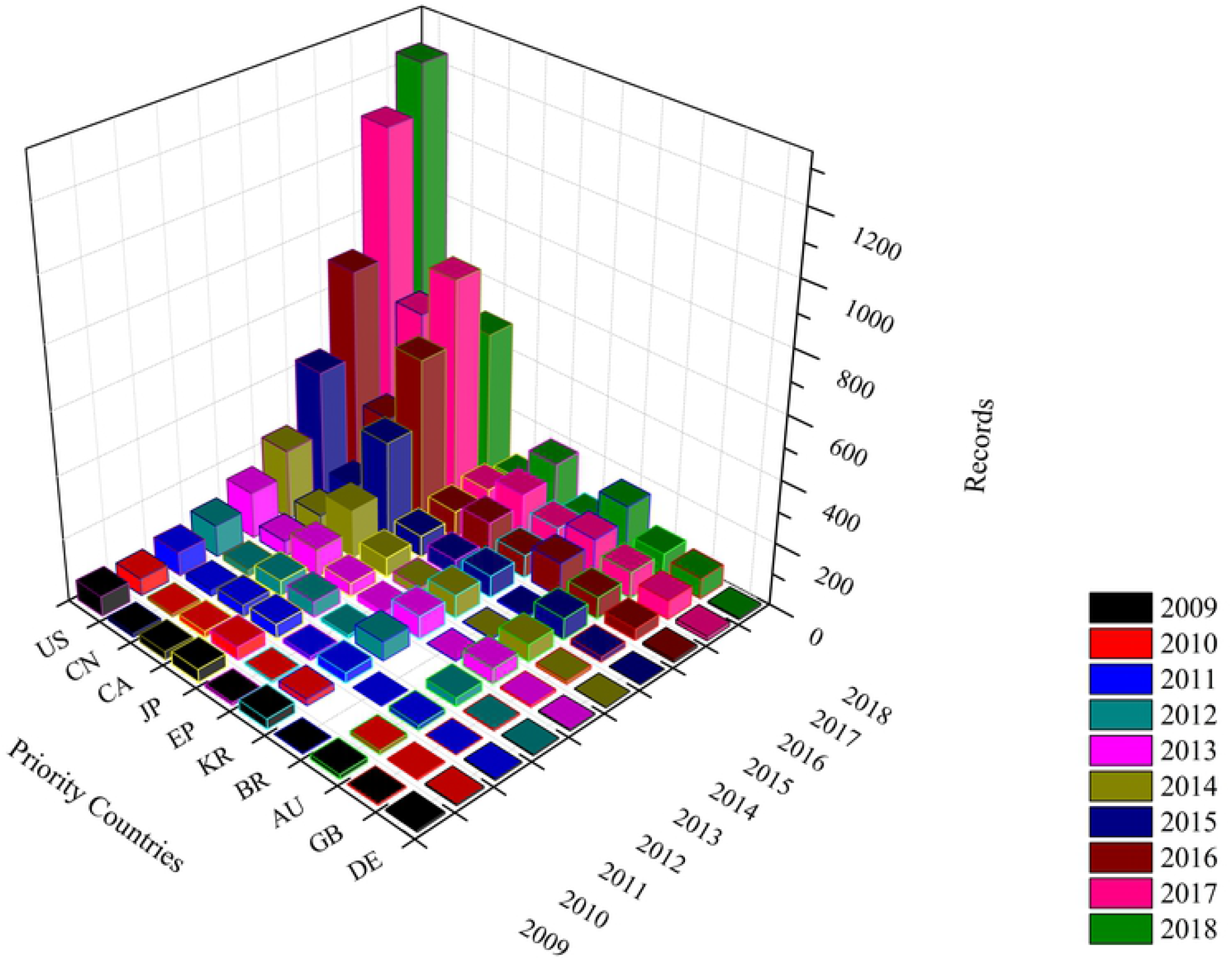
The annual changing trends of the top 10 priority countries in the field of antineoplastics targeting PD-1/PD-L1 from 2009 to 2018.

#### Patent Assignees

The top 15 patent assignees were shown in Fig 9. More than half are American patent assignees, with 8 ones, including three companies, two research institutes, two universities, one hospital and one government department. Genentech was an American biotechnology company, and became a subsidiary of Roche in 2009. Others were three Japanese companies, two Swiss companies, one Korean company and one French institute. The top three patent assignees were Bristol-Myers Squibb (USA), Dana-Farber Cancer Institute (USA), and Novartis (Switzerland). China ranked second place in the number of patent applications in priority countries, but no Chinese patent assignees ranked among the top 15 in the world.

**Fig 9.**
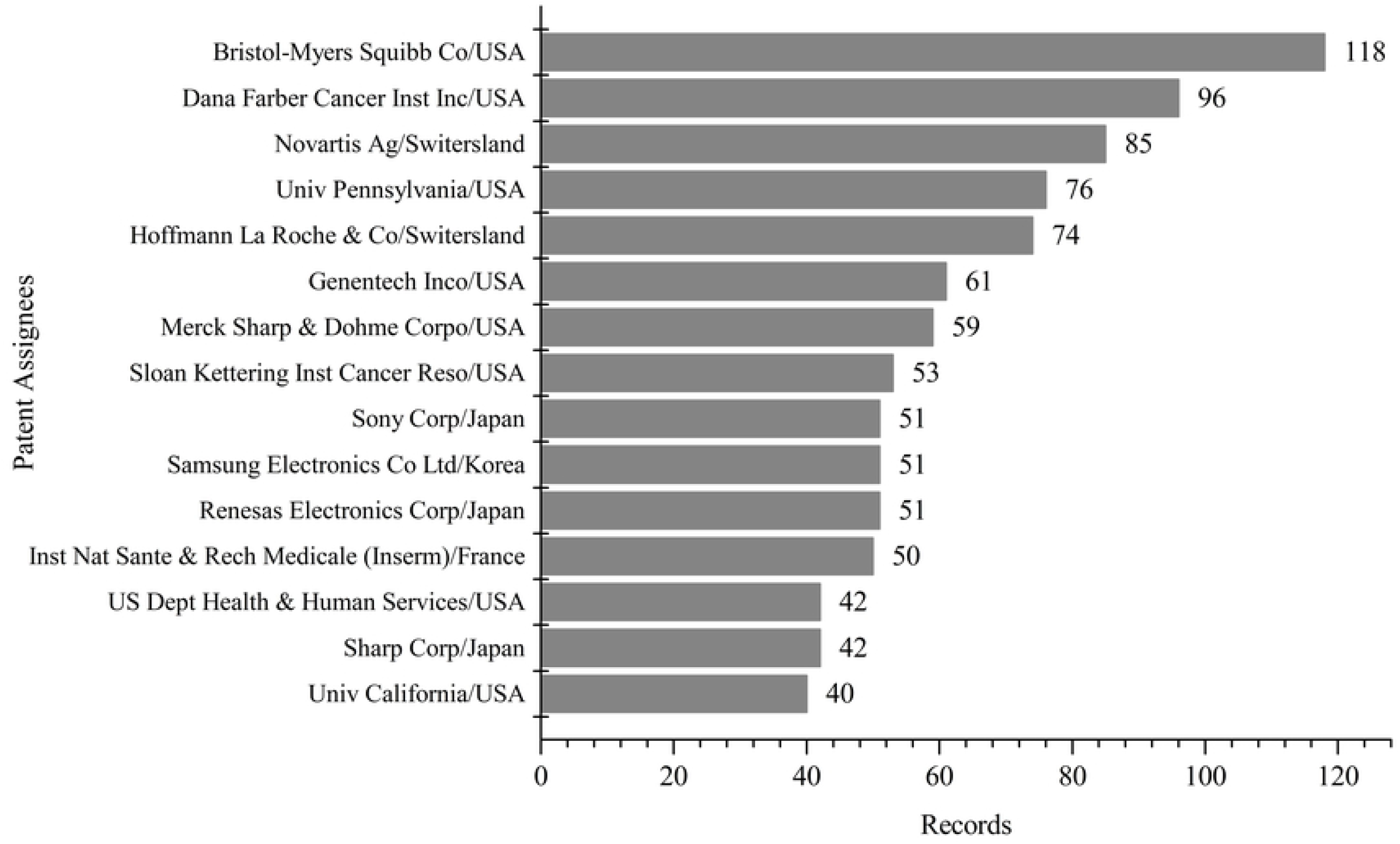
The top 15 patent assignees in the field of antineoplastics targeting PD-1/PD-L1.

#### Technological Development Focus

There were 5501 patents in the field of antineoplastics targeting PD-1/PD-L1, the titles and abstracts of which were chosen to identify the technological development focus, with the same method and procedure of identifying the scientific research focus, which were also imported to VOSviewer for clustering and visualization. Fig 10 showed the clusters of co-word matrix made by VOSviewer, and there were divided into five clusters, one more focus than the scientific research ones, indicating that the technological development focus of related to antineoplastics targeting PD-1/PD-L1 were more diverse. The five technological development focus were as follows: (1) testing methods and apparatus, (2) indications related to carcinoma, (3) biomarkers related to diagnosis and prognosis, (4) small molecule inhibitors, and (5) indications other than carcinoma (Fig 10A). Those words associated with each focus were listed in Table 2, and each cluster only listed the top ten high frequency words due to the limited space. Fig 10B showed the technological development focus related to antineoplastics targeting PD-1/PD-L1, which indicated that the five focus were distributed from 2008 and 2016, and three focus of indications related to carcinoma, biomarkers related to diagnosis and prognosis, and small molecule inhibitors were newer, the average application years of which were all almost in 2016. And the focus of indications other than carcinoma was a little older than the above three ones, with the average application year of 2014 nearly. Then, the focus of testing methods and apparatus was the oldest, with the average application year close to 2008. So, as could be seen from the Fig 10B, the technological development focus related to antineoplastics targeting PD-1/PD-L1 was on testing methods and apparatus initially, and then had changed over time. The three focus of indications related to carcinoma, biomarkers related to diagnosis and prognosis, and design and synthesis of derivatives had received more attention in recent years.

**Table 2.**
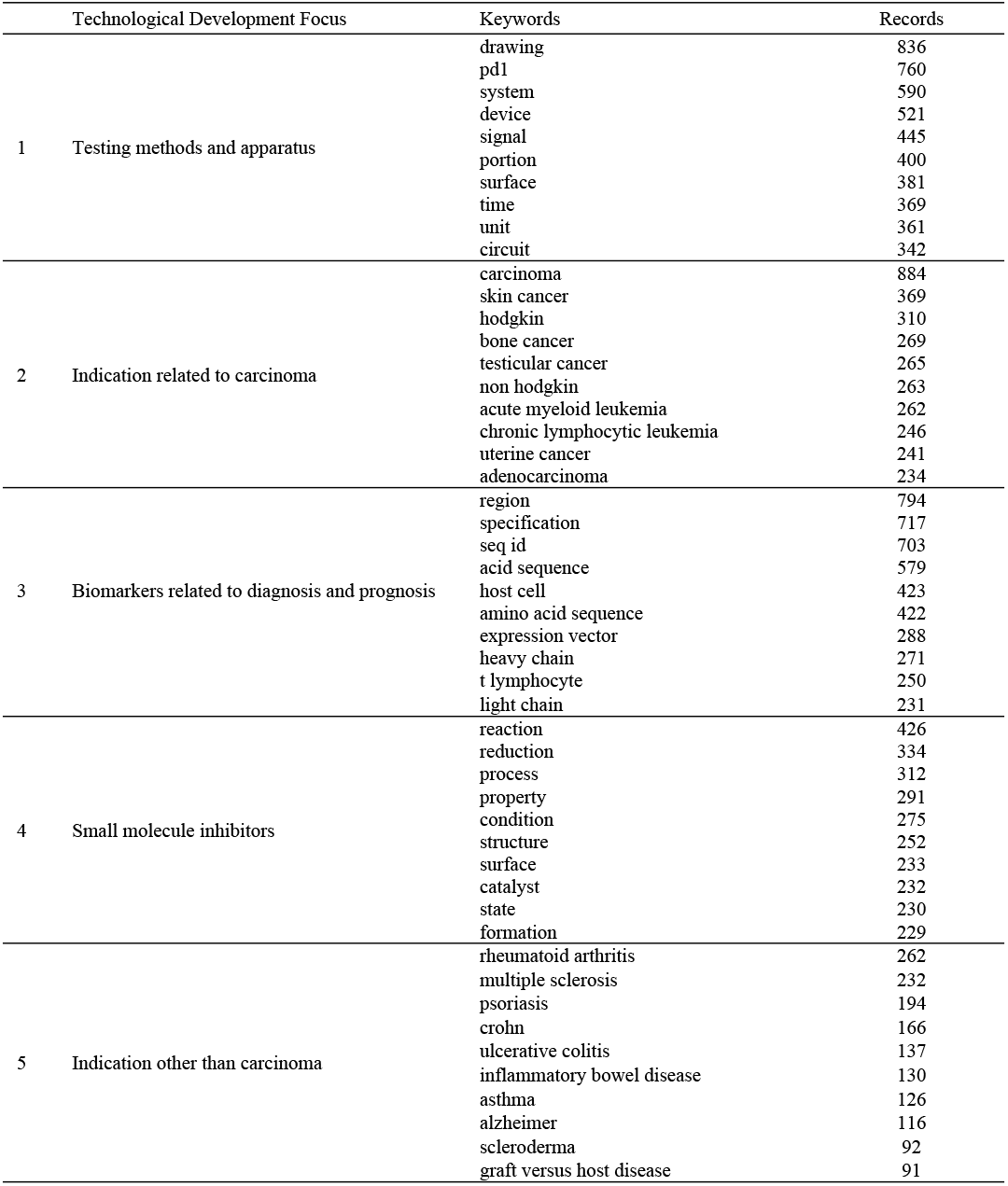
The technological development focus related to antineoplastics targeting PD-1/PD-L1.

**Fig 10.**
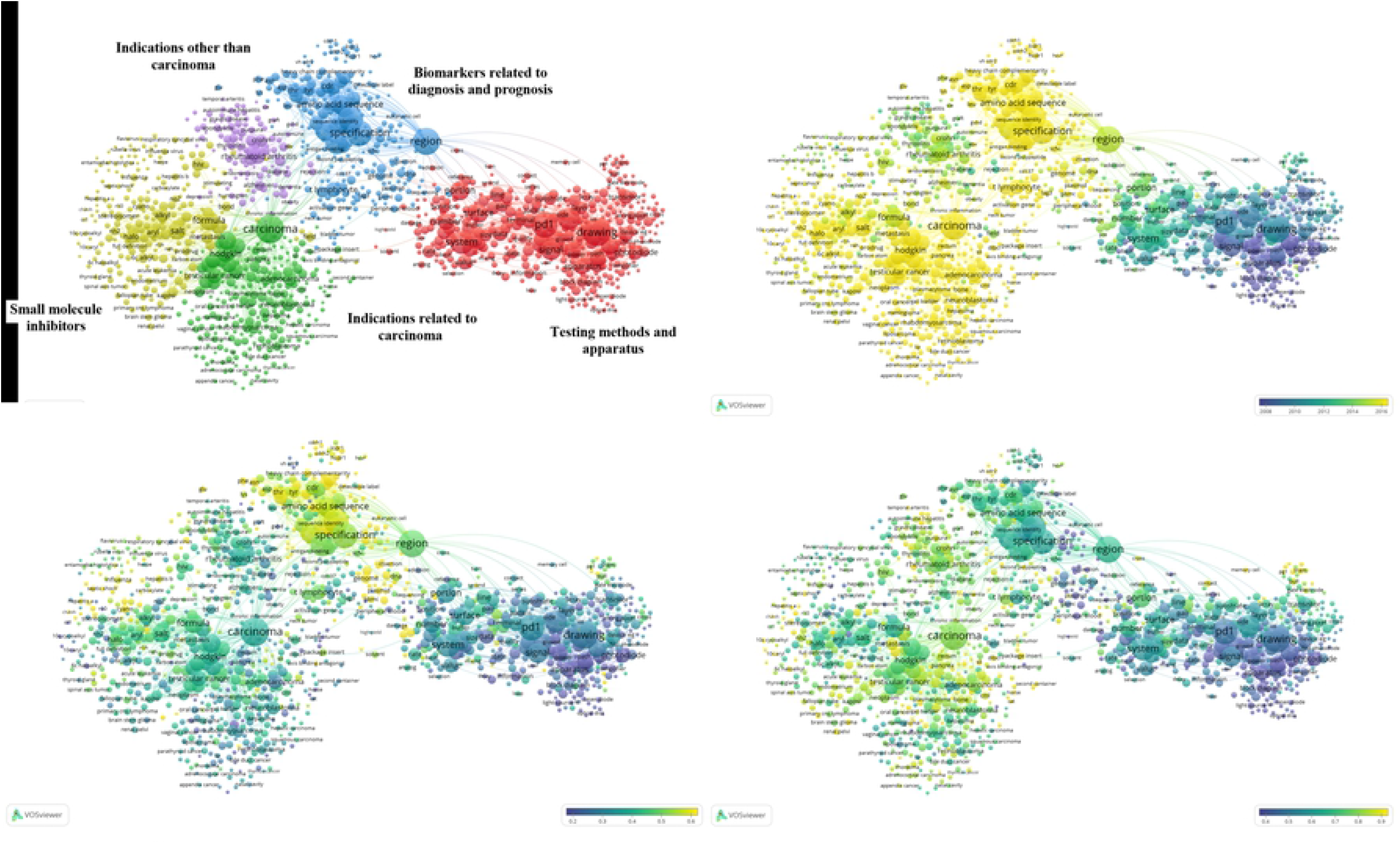
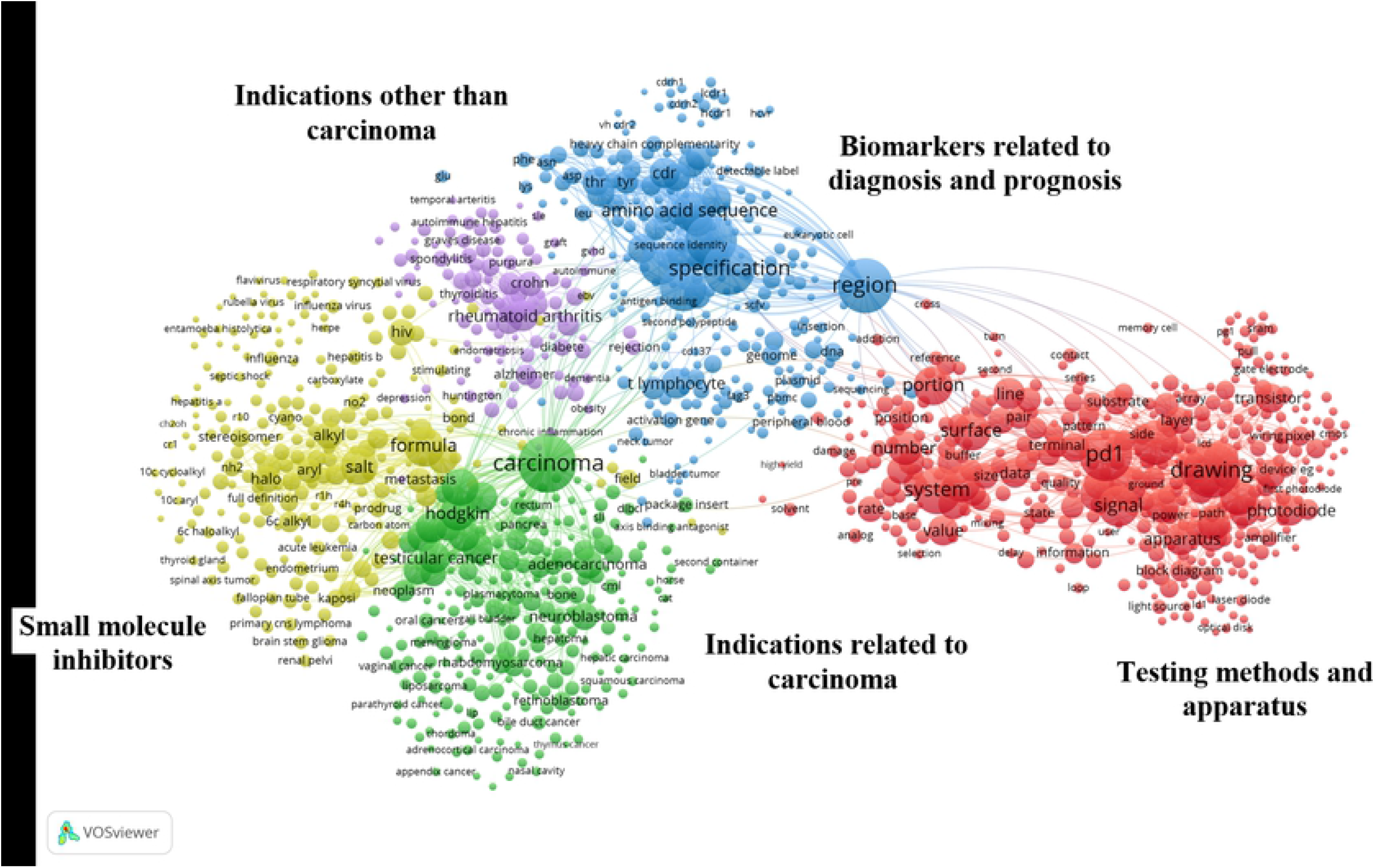
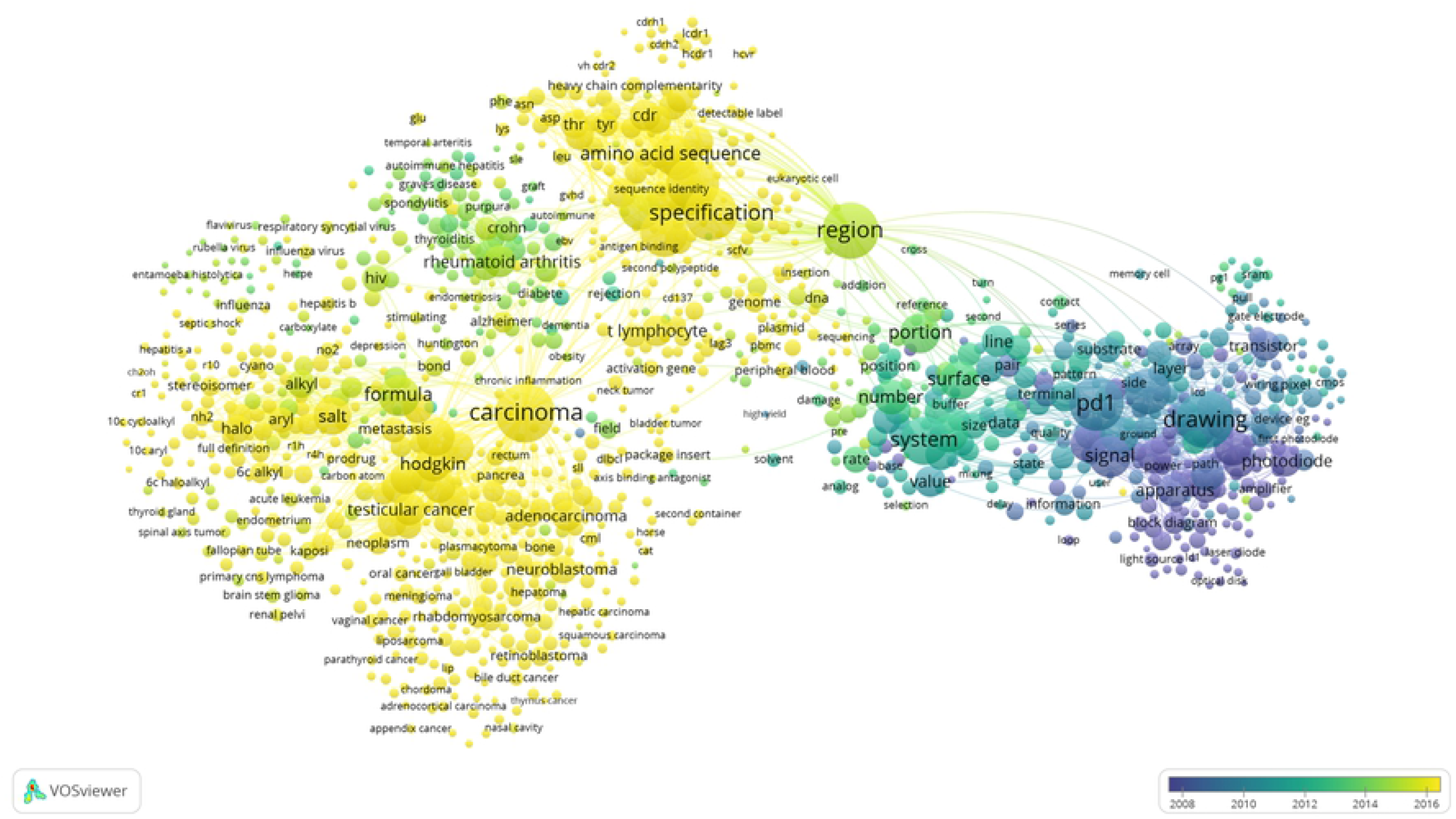
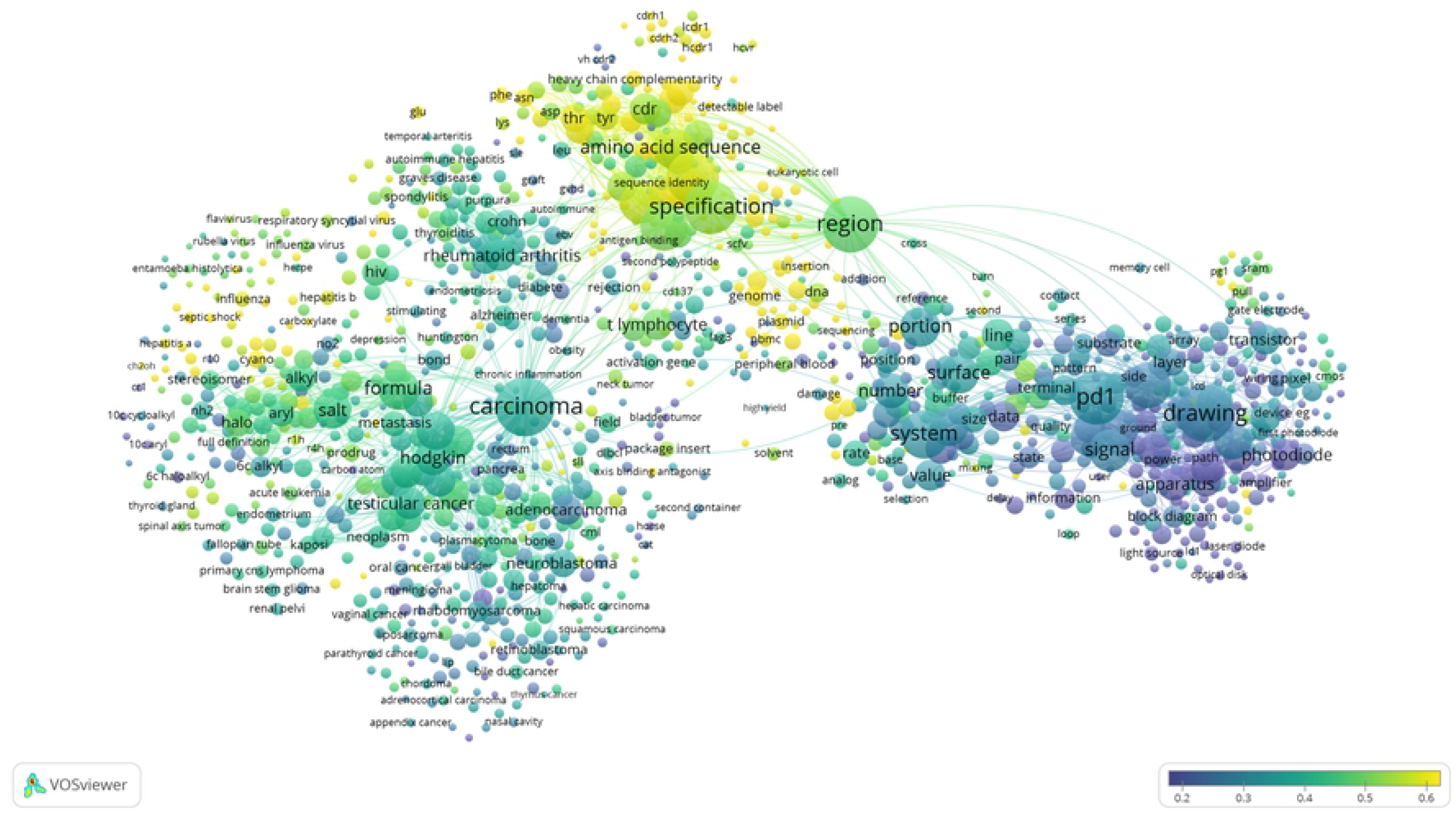
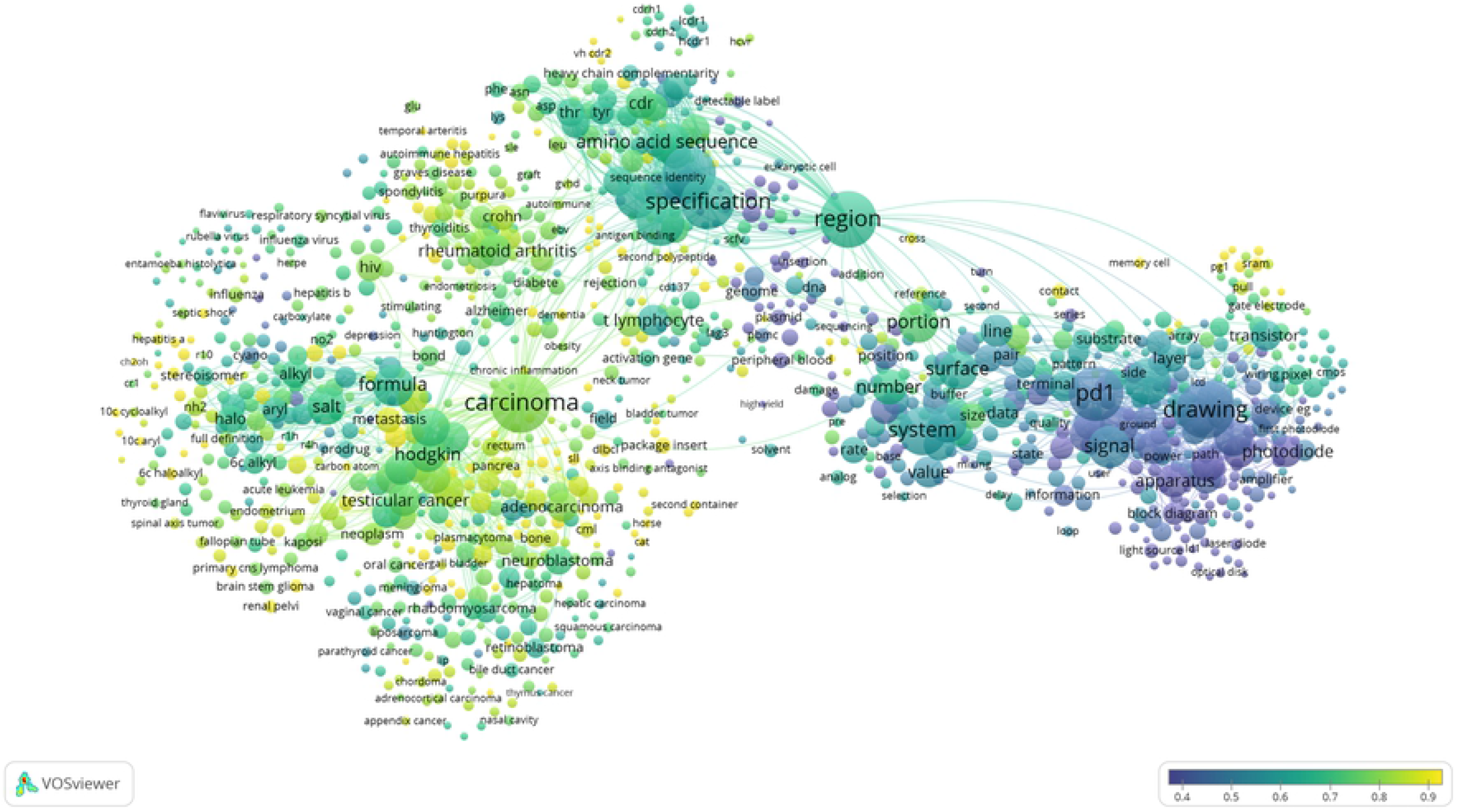
The technological development focus related to antineoplastics targeting PD-1/PD-L1. **(A)** The visualization of the technological development focus related to antineoplastics targeting PD-1/PD-L1 based on the VOSviewer. **(B)** The temporal development of the technological development focus related to antineoplastics targeting PD-1/PD-L1in recent five years. The color is from purple to yellow. The closer to purple (the year 2008), the focus is older, and the closer to yellow (the year 2016), the focus is newer. **(C)** The technological development focus of China related to antineoplastics targeting PD-1/PD-L1. Colors from purple to yellow indicate that the concern of focus is low (0.2) to high (0.6). The color is from purple to yellow. The closer to the purple (0.2), the focus is lower concerned, and the closer to the yellow (0.6), the focus is higher concerned. **(D)** The technological development focus of USA related to antineoplastics targeting PD-1/PD-L1. Colors from purple to yellow indicate that the concern of focus is low to high. The color is from purple to yellow. The closer to the purple (0.4), the focus is lower concerned, and the closer to the yellow (0.9), the focus is higher concerned.

##### Indications related to carcinoma

This is the way to expand the indications. The earliest PD-1 / PD-L1 inhibitors used in clinical trials were pembrolizumab and nivolumab, both of which were initially approved for the treatment of melanoma ^[47,48]^. Subsequently, pembrolizumab was approved for the treatment of non-small cell lung cancer ^[49]^, and nivolumab was approved for the treatment of renal cell carcinoma ^[50]^, non-small cell lung cancer ^[51]^, and squamous-cell carcinoma of the head and neck ^[52]^. It is a fast and effective way to find antineoplastics by the way of constantly exploring the therapeutic efficacy of existing tumor immunotherapy drugs in other tumors. The discovery of new uses is an important direction for new drug discovery, and this is drug repositioning ^[53]^. The new drugs discovery through new uses of existing drugs has significantly reduced the cost of research and development, expanded the indications and clinical treatment scope of the original drugs, and the clinical application has tended to be safe, reasonable and effective, with reducing the risk of discovering new drugs in the traditional way at the same time ^[54]^. Drug repositioning strategy is one of the best risk-benefit ratio strategies in the currently known drug discovery strategies, and it is also one of the effective methods to solve the dilemma of high investment and low success rate in new drug discovery ^[55–57]^. The rapid development of tumor immunotherapy and the promising clinical research results have prompted the continuous development of immune checkpoint inhibitors, which are actively researched and expanded to many other types of malignant tumors, and they need clinical trials to be approved for marketing. Although most of these results of are in further clinical trials, this kind of new drugs are rapidly changing the landscape of cancer treatment.

##### Biomarkers related to diagnosis and prognosis

This technological development focus is similar to the scientific research focus of biomarkers related to efficacy and prognosis. With no doubt, advances in scientific research have also contributed to the technological development on biomarkers. More and more research institutions and enterprises have accelerated the protection of biomarker products in the field of intellectual property. These patents are mainly the methods of tumor diagnosis or prognosis by biomarkers, including the new biomarkers discovery, new or improved diagnostic methods, etc ^[58–62]^. The research results of biomarkers for diagnosis and treatment can directly guide the development of diagnostic tools and the selection of therapeutic drugs, which have great market prospects. Developing reliable biomarkers for predicting efficacy and prognosis is one of the major challenges in the field of tumor immunotherapy, and it will remain the focus of scientific research and technological development ^[63]^. Scientific research is the foundation and source of technological development, and also technological development will promote scientific research in turn. The interactive relationship between science and technology can promote the development of biomarkers better and faster in the future.

##### Small molecule inhibitors

Although antibody-based tumor immunotherapy has significant clinical efficacy and has made many breakthroughs, there are still some unavoidable problems. The monoclonal antibodies can cause serious immune-related adverse events (irAEs), due to the long half-life and the binding to targets for too long ^[64,65]^. In addition, the production process of monoclonal antibodies is complicated, expensive and not easy to store and transport. Compared with monoclonal antibodies, small molecule drugs have many significant advantages, such as low price, oral administration, easy transportation and storage, and non-immunogenicity ^[66,67]^. Therefore, searching for small molecule inhibitors of PD-1 / PD-L1 has become a hot spot for new drug discovery. At present, the reported small molecule inhibitors include the following types: peptides ^[68,69]^, sulfonamides ^[70]^, biphenyls ^[71,72]^, oxadiazoles ^[73]^, etc. The first small molecule inhibitor of PD-1 was a class of sulfonamides discovered by Harvard University in 2011, and such sulfonamide derivatives were expected as lead compounds to be further modified and optimized to design compounds with druggability ^[70]^. Comparing with antibody drugs, the binding activity of small molecule inhibitors to targets is lower. However, small molecule inhibitors have controllable pharmacokinetic properties and mature research system, which makes it possible to overcome the problems of antibody drugs. Therefore, it is very important to design and synthesize small molecule inhibitors that block the binding of PD-1 / PD-L1.

Fig 10C and Fig 10D showed the different technological development focus between China and USA. It could be seen from the Fig 10C that the focus of biomarkers related to diagnosis and prognosi (between 0.5 and 0.6) was most concerned in China. The three focus of indications other than carcinoma, small molecule inhibitors, and indications related to carcinoma were less concerned. It seemed that the focus of testing methods and apparatus received the least attention in China, comparatively. The American focus were a little different, and there were three most concerned focus (between 0.7 and 0.9), such as indications other than carcinoma, small molecule inhibitors, and indications related to carcinoma. The focus of biomarkers related to diagnosis and prognosis was less concerned in USA (between 0.6 and 0.7), while it received most attention in China. As was the same in China, the focus of testing methods and apparatus received the least attention in USA.

## Conclusion

This study uses scientometrics and patentometrics to explore the research activity and reveal the current characteristics in the field of antineoplastics targeting PD-1/PD-L1. This field has received widespread attention, growing rapidly in recent years. It is predicted that the document publications and the patent applications will reach 7596 and 6553 in 2023, severally. USA is the most productive country, not only with the greatest share of publications, but also the largest number of patents. Also, the top one publishing organization and top one patent assignee are also from USA, and they are Harvard University and Bristol-Myers Squibb, respectively. China ranks second in the world, but there is a large gap in both the number of document publications and the number of patent applications compared with USA.

## Acknowledgments

This study is supported by the Fundamental Research Funds for the Central Universities (NO. 2017PT63006) and the National Key Research and Development Program of China (NO. 2016YFC0104805).

## Author Contributions

TZ conceived and designed the experiments. TZ performed the experiments. TZ, JC and YL analyzed the data. TZ contributed materials and analysis tools. TZ wrote the paper. TZ, ZLOY obtained the funding. TZ has primary responsibility for the final content.

## Declaration of interest

All authors read and approved the final manuscript. The authors have declared that no competing interests exist.

## Notes

**Funding:** This study is supported by the Fundamental Research Funds for the Central Universities (NO. 2017PT63006) and the National Key Research and Development Program of China (NO. 2016YFC0104805).

